# Glycan-Protein Interactions Determine Kinetics of *N*-Glycan Remodeling

**DOI:** 10.1101/2020.12.01.406371

**Authors:** Corina Mathew, R. Gregor Weiß, Christoph Giese, Chia-wei Lin, Marie-Estelle Losfeld, Rudi Glockshuber, Sereina Riniker, Markus Aebi

## Abstract

A hallmark of *N*-linked glycosylation in the secretory compartments of eukaryotic cells is the sequential remodeling of an initially uniform oligosaccharide to a site-specific, heterogeneous ensemble of glycostructures on mature proteins. To understand site-specific processing, we used protein disulfide isomerase (PDI), a model protein with five glycosylation sites, for molecular dynamics (MD) simulations and compared the result to a biochemical *in vitro* analysis with four different glycan processing enzymes. As predicted by an analysis of the accessibility of the *N-*glycans for their processing enzymes derived from the MD simulations, *N*-glycans at different glycosylation sites showed different kinetic properties for the processing enzymes. In addition, altering the tertiary structure context of *N*-glycan substrates affected *N*-glycan remodeling in a site-specific way. We propose that differential, tertiary structure context dependent *N*-glycan reactivities lead to different glycan structures in the same protein through kinetically controlled processing pathways.

## Introduction

*N*-glycoprotein biogenesis in eukaryotes is initiated in the Endoplasmic Reticulum (ER) by the oligosaccharyltransferase, an enzyme complex which covalently links a uniquely defined oligosaccharide, Glc_3_Man_9_GlcNAc_2_, to the side-chain amide nitrogen atom of asparagines within −N-X-S/T-sequons. It thereby modifies a large number of proteins, many of them at multiple sites [1–4]. After the transfer to the protein, processing of the *N*-linked glycans is initiated by ER-localized hydrolases. The removal of the three glucoses is hereby coupled to the folding of the glycoproteins by providing ligands for lectin chaperones such as calnexin or calreticulin [5, 6]. The quality control of protein folding relies on glycans and the information they provide about the conformational state of the covalently bound protein [7].

After the exit from the ER, *N*-glycoproteins are further processed by Golgi specific hydrolases and transferases that generate the final structures of *N*-glycans [8]. This remodeling pathway is characterized by individual reactions that rarely go to completion and the processing of a glycan being different for each site of the glycoproteome. Consequently, a site-specific heterogeneity of *N*-glycan structures is observed. Early on, it was suggested that this differential processing might be due to the tertiary structure of the glycoprotein. In 1984, Savvidou *et al.* hypothesized that the decreased amount of bisecting *N*-glycans on one specific glycosylation site of human IgG was due to specific interactions between this glycan and the protein [9]. In the following, several NMR studies demonstrated interactions between *N*-glycans and the protein surface, which are mostly facilitated by the reducing-end GlcNAc of an *N*-glycan [10, 11]. A most recent example of tertiary structure context dependent glycan processing is the discovery that only a single of the eight N-glycans in the filamentous urinary glycoprotein uromodulin (UMOD) remains a high-mannose type glycan, while all other *N*-glycans are further processed to complex-type *N*-glycans. It is however the single high-mannose type UMOD glycan that mediates encapsulation and aggregation of uropathogens by UMOD filaments via interactions with mannoside-specific pilus lectins from the pathogens [12]. In addition, the rise of computational glycobiology allowed the simulation of glycan-protein interactions [13, 14] and indicated that these interactions could reduce the accessibility of the glycan to glycan-processing enzymes [15]. Consequently, glycan-protein interactions are considered a major determinant of *N*-glycan microheterogeneity [15, 16]. Several studies have used this knowledge to engineer glycoproteins by site-directed mutagenesis. Chen *et al.* introduced new glycan-protein interactions into IgG and could thereby significantly improve the stability of IgG against thermal and low pH induced aggregation [17]. In contrast, site-directed amino acid replacements disrupting interactions between the glycan and the protein lead to improved processing of the “freed” *N*-glycans [18, 19].

Understanding site-specific *N*-glycan processing as it occurs *in vivo* requires a detailed knowledge of the specificity and localization of the individual hydrolases and glycosyltransferases [20]. After the removal of the three terminal glucoses in the ER, the α-1, 2-mannosidase ER mannosidase I (ER Man I) removes the terminal mannose from the B-branch of the *N*-glycan. Even though ER Man I has a high specificity towards this terminal mannose, it is capable of trimming all α-1, 2-linked mannoses from an *N*-glycan [21, 22]. In the Golgi, the glycan is further processed by Golgi Mannosidase I (GM I). Belonging to the same glycoside hydrolase family as ER Man I, this enzyme is also able to remove all α-1, 2-linked mannoses from the *N*-glycan [23]. When confronted with a Man_9_GlcNAc_2_ glycan, GM I works least efficiently on the terminal B-branch mannose, the preferred mannose of ER Man I [24].

The resulting Man_5_GlcNAc_2_ glycan is further processed by *N*-acetylglucosaminyltransferase I (GnT I), which transfers one GlcNAc to the A-branch of the glycan and uses UDP-GlcNAc as a donor substrate [25]. Its action is essential, as the transfer of a GlcNAc initiates the formation of hybrid *N*-glycans [26]. The generated GlcNAcMan_5_GlcNAc_2_ serves as a substrate for Golgi mannosidase II (GM II) which can cleave two mannoses with different glycosidic linkages (α-1, 3 and α-1, 6 linked), of the B- and the C-branch, with a single catalytic site. First, the α-1, 6-linked, terminal mannose is removed, followed by the α-1, 3 linked mannose [27, 28]. By the removal of these two mannoses GM II initiates the synthesis of complex glycan structures [29].

We used yeast protein disulfide isomerase (PDI) as a model protein with five *N*-glycosylation sites to investigate site-specific *N*-glycan processing in the context of an intact glycoprotein [15]. We performed in-depth molecular dynamics (MD) simulations to analyze the dynamics and the interactions of the *N*-linked glycans and experimentally addressed site-specific processing by ER Man I, GM I, GnT I and GM II *in vitro.* Initial velocities and *K*_*M*_ values demonstrated that the glycan of each glycosylation site represents a unique substrate to glycan-processing enzymes. MD simulations explained the site-specific properties that were primarily determined by protein-glycan and glycan-glycan interactions. Altering the protein structure changed site-specific glycan processing, validating the conclusion that intramolecular protein/glycan interactions slow or even prevent individual steps of glycan processing.

## Results

### Molecular dynamics simulations of PDI

Figure 1A schematically illustrates the chemical composition of a Man_9_GlcNAc_2_ glycan, the most abundant structure bound to the glycosylation sites of PDI in the ER. The U-shaped PDI structure shown in figure 1B consists of four thioredoxin-like domains termed a, b, b’, and a’, of which the domains a and a’ possess a catalytic cysteine pair (CGHC in figure 1B) [30]. Additionally, the a-domain contains a structural disulfide bond (C-C in figure 1B). The glycosylation sites 1 and 2 are located on the a-domain, while the b-domain contains sites 3 and 4. Site 5 is distantly located from the other glycosylation sites on the a’-domain.

**Figure 1:**
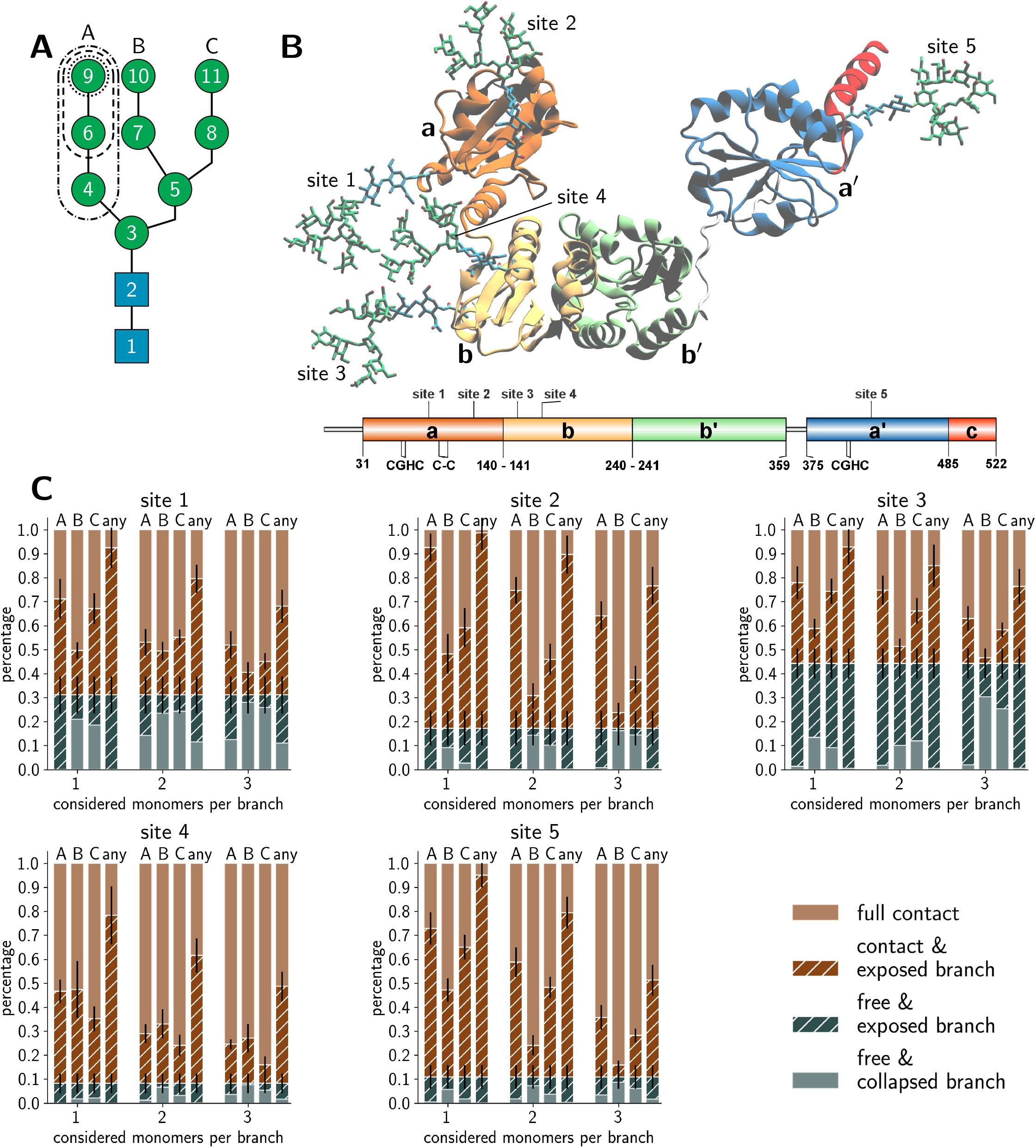
Structural assessment of the site-specific glycan accessibility. **A**: Simplified representation of a branched Man_9_GlcNAc_2_ oligosaccharide. The schematic also exemplifies on the A-branch the considered branch lengths for the accessibility assessment when accounting for one (dotted line), two (dashed line), or three monomers (dotted-dashed line). **B**: Structure of a Man_9_GlcNAc_2_ glycoform of PDI. The secondary structure of the protein is shown as ribbon representation, the glycans are in a licorice representation with mannose residues in green and *N*-acetylglucosamine residues in blue. The four thioredoxin domains are labeled in different colors (a, orange; b, yellow; b’, green; a’, blue). The sequence of PDI indicates the three disulfide bonds. The first disulfide bond on the a domain and the one on the a’ domain are the active sites of PDI (CGHC). The second disulfide bond on the a domain (C-C) is structural. The glycosite positions are also indicated. **C:** Accessibility analysis of the Man_9_GlcNAc_2_ glycans of PDI. The bar plots show the sum of probabilities from the stationary distribution of microstates that are classified as ‘free’ (blue) and in ‘contact’ (brown). The striped color refers to ‘free’ and ‘contact’ states that have either the A, B, C, or any branch exposed. The abscissa labels the classifications based on one, two, or three branch monomers.

We simulated an aggregate sampling of 75 μs for the full-length glycoprotein with glycosylated sites 1–4 and 110 μs for the a’-domain with glycosylated site 5. The MD simulations were partitioned into structural microstates and analyzed using Markov state modeling (MSM) [31, 32], typically used to reconstruct thermodynamic and kinetic properties of a simulated ensemble. From the MSM, we retrieved the stationary probability distribution of the structural glycan microstates to quantify the amount of accessible glycan conformations. Further details on the MD simulations, microstate definition, MSM construction, and accessibility assessment are given in the Methods section.

### Conformations of PDI glycans

We distinguished four categories in total for the branch accessibility. The two categories with ‘free’ glycan classification with or without exposed branch(es) were labeled as ‘free & exposed branch’ and ‘free & collapsed branch’, respectively. The other two categories differentiate glycan conformations in ‘contact’ with the protein environment but with or without branch exposure, respectively labeled as ‘contact & exposed branch’ and ‘full contact’. Thus, the classification ‘exposed’ indicated glycan structures, for which a branch was stretching into the solvent while the others could still be interacting with the protein, neighboring glycans, or other branches within the same glycan. From the enzymatic point of view, extended glycan conformations are preferred because the catalytic sites in GH47 α-mannosidases are known to be deep funnels binding only one extended branch at a time [33]. Hence, the ‘contact & exposed branch’ conformations could be more beneficial for the deep binding funnels than the ‘free & collapsed branch’ conformations.

The accessibility assessment of sites 1–5 is shown in figure 1C. Sites 1 and 3 exhibited the largest populations of ‘free’ conformations in which the glycan was barely interacting with its surrounding protein environment or neighboring glycans. Site 2 also showed a clear but reduced fraction of ‘free’ conformations but had a considerable amount of ‘exposed’ microstates. Interestingly, a common pattern of ‘exposed’ branches was shared among sites 1–3 and 5. The ‘exposed’ microstates were dominated by ‘exposed’ A-branch conformations, the most flexible branch. A generally minimal exposure of the B branch is related to its central location within the glycan. Naturally, the percentages of exposed conformations declined when considering an increasing number of monosaccharides per branch. This gradient was site-specific. In contrast, the same assessment of the glycan on site 4 showed a significantly different pattern. The percentage of ‘full contact’ conformations on site 4 was most dominant across all levels of branch lengths, while the fraction of ‘free’ conformations was the lowest compared to all other sites. Furthermore, the amount of conformations in which a single branch is completely solvent exposed (i.e. ‘free & exposed’ conformations) was reduced. In contrast, the preference of A and C branch exposure over the B branch was lost, and all three branches had similar fractions of ‘exposed’ conformations. In summary, our glycan-centric, quantitative analysis of the microstates of the MSMs suggested that the Man_9_GlcNAc_2_ was least accessible on site 4, had a tendency to branch exposure but contacting conformations on sites 2 and 5, while sites 1 and 3 exhibited patterns with large solvent exposure. These observations indicated site-specific differences between the five (Man_9_GlcNAc_2_) sites in their reactivity with ER Man I despite their identical chemical structures.

The above analysis of individual glycan conformations was not accounting for the particular contacts and interactions with the glycans’ environments that would lead to the ‘free’, ‘contact’, and ‘exposed’ classifications. For instance, as illustrated in figure 1B, sites 1 and 3 are in close proximity to each other such that glycan-glycan interactions contributed dominantly to the fraction of ‘contact’ conformations. Hence, a competition of site 1 and 3 during glycan-enzyme interaction could reduce the trimming of the branches on either site. Figure 1B shows further that the glycan on site 5 can easily extend to free conformations but potentially forms frequent interactions with the acidic, C-terminal α-helical PDI segment.

Thus, while the amount of free conformations was clearly affected by the glycan–protein interactions at site 5, the individual branches are still exposed during the contact with the neighboring α-helix. In addition, the convex and concave surrounding protein surface topology at site 2 and 4, respectively, pose different glycan–protein contact possibilities (figure 1B and figure EV1). At site 2, the surrounding protein surface has a positive curvature such that the branches are easily extended when the glycan is ‘free’ or in ‘contact’ (figure EV1). In contrast, the concave protein surface around site 4 hinders ‘exposed’ branches in the ‘contact’ and ‘free’ conformations.

### *In vitro* processing of PDI *N*-glycans

#### Experimental setup

To experimentally test the predictions offered by the MD simulations, we turned to an *in vitro* biochemical analysis of *N*-glycoprotein processing. We therefore analyzed the kinetics of *N*-glycan maturation processes of the ER and Golgi (figure EV2A) by incubating PDI with the *N*-glycan processing enzymes ER Man I, GM I, GnT I, and GM II. PDI was prepared for (mainly) uniform Man_9_GlcNAc_2_, Man_5_GlcNAc_2_, or GlcNAcMan_5_GlcNAc_2_ glycosylation corresponding to the respective enzyme’s glycan specificity (figure EV3).

After incubation of glycosylated PDI with the respective remodeling enzyme, the site-specific glycoform distributions were obtained by mass spectrometry (MS) of tryptic glycopeptides (figure EV2B).

#### Site-specific processing of *N*-glycans by different enzymes

For processing of Man_9_GlcNAc_2_-PDI (60 μM) by ER Man I (0.3 μM), we observed efficient conversion (90%) from Man_9_GlcNAc_2_ to Man_8_GlcNAc_2_ for sites 1-3 and 5 within approximately three minutes. In contrast, the glycan from glycosite 4 was processed significantly slower. After 60 minutes only 85% conversion was obtained. On site 2, we additionally observed a decrease of the Man_8_GlcNAc_2_ product after 10 minutes, corresponding to additional mannose trimming by ER Man I (figure 2A).

**Figure 2:**
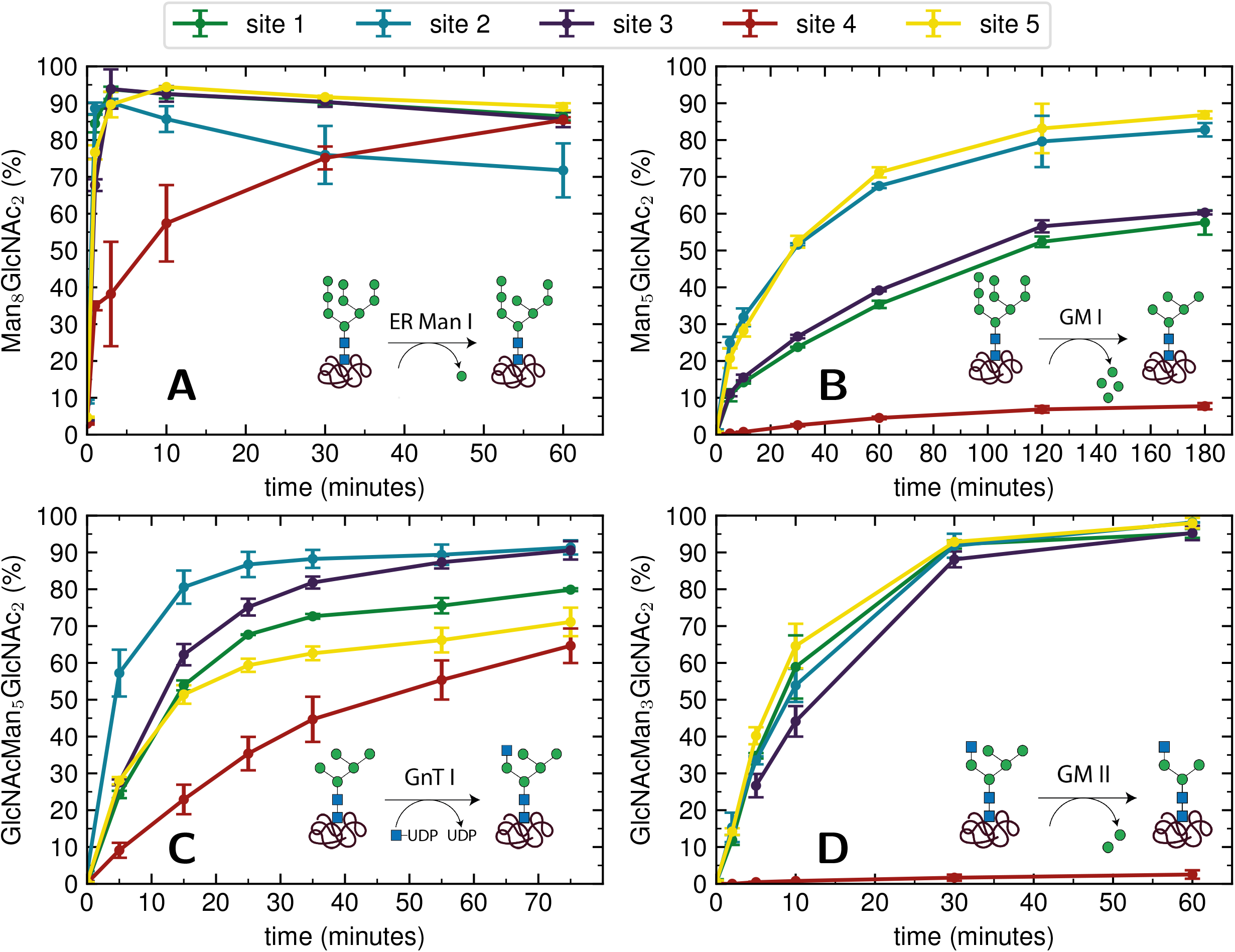
Site-specific processing of PDI *N*-glycans by four different enzymes. **A:** Reaction catalyzed by ER Man I: Removal of one α-1, 2-linked mannose from the B-branch of Man GlcNAc, resulting in Man GlcNAc . Man GlcNAc -PDI was incubated with purified ER Man I, shown is the conversion of Man GlcNAc to Man GlcNAc on each site over 60 minutes. Error bars represent the standard deviation of three independent experiments (n=3). **B:** Reaction catalyzed by GM I: Removal of four α-1, 2-linked mannoses from Man_9_GlcNAc_2_, resulting in Man_5_GlcNAc_2_. Man_9_GlcNAc_2_-PDI was incubated with GM I, shown is the conversion of Man_9_GlcNAc_2_ to Man_5_GlcNAc_2_ on each site over 180 minutes (*n*=3). **C:** Reaction catalyzed by GnT I: Transfer of a GlcNAc from UDP-GlcNAc to the A-branch of a Man_5_GlcNAc_2_ glycan, resulting in GlcNAcMan_5_GlcNAc_2_. Man_5_GlcNAc_2_-PDI was incubated with purified GnT I and UDP-GlcNAc. Shown is the conversion of Man_5_GlcNAc_2_ to GlcNAcMan_5_GlcNAc_2_ over 80 minutes (*n*=3). **D:** Reaction catalyzed by GM II: Removal of a α-1, 3-linked mannose from the B-branch and a α-1, 6-linked mannose from the C-branch of a GlcNAcMan_5_GlcNAc_2_ glycan, resulting in GlcNAcMan_3_GlcNAc_2_. GlcNAcMan_5_GlcNAc_2_-PDI was incubated with purified GM II and shown is the conversion of GlcNAcMan_5_GlcNAc_2_ to GlcNAcMan_3_GlcNAc_2_ on each site over 60 minutes (*n*=3).

For processing of Man_9_GlcNAc_2_-PDI (20 μM) by GM I (0.1 μM), site 4 again proved to be processed slowest: Only 8% of the site 4 glycans were converted to Man_5_GlcNAc_2_ (in contrast to 80% at sites 2 and 5, figure 2B).

The processing of Man_5_GlcNAc_2_ by glycosyltransferase GnT I also proved to be slowest at site 4. After incubation of Man_5_GlcNAc_2_-PDI (30 μM) with GnT I (0.15 μM) and UDP-GlcNAc (5 mM) only about 65% of the site 4 glycans were converted to GlcNAcMan_5_GlcNAc_2_, while glycans at sites 2 and 3 were converted to about 90% (figure 2C).

The largest differences in site-specific remodeling could be observed for GM II (figure 2D). Sites 1-3 and 5 of GlcNAcMan_5_GlcNAc_2_-PDI (20 μM) reacted to more than 90% to GlcNAcMan_3_GlcNAc_2_ in the presence of GM II (67 nM) within 60 minutes, while hardly any conversion could be observed on site 4.

Hence, we observed that for all four enzymes, despite being hydrolases or transferases, site 4 was processed slowest.

#### Michaelis-Menten analysis of processing kinetics

To quantify the site-specific differences in glycan processing kinetics for the one-step reactions catalyzed by ER Man I and GnT I, we next recorded the dependence of the initial velocity of substrate conversion on PDI concentration at constant processing enzyme concentration (ER Man I: 0.6 nM; GnT I: 67 nM). Site-specific differences in glycosylation site occupancy of PDI (table EV3) and efficiency of kifunensine treatment (for the ER Man I assay) or GM I treatment (for the GnT I assay) were analyzed by MS in order to calculate the actual concentration of substrate glycan structure per site.

Initial velocities of ER Man I (figure 3A) and GnT I (figure 3B) for all five sites against PDI concentration were plotted together with fits according to a Michaelis-Menten mechanism. We observed initial velocities increased with PDI concentrations towards saturation (table EV4–5, figure EV4). While the initial velocities of GnT I approached *v*_*max*_ for all sites, they still increased nearly linearly on site 4 for ER Man I. Consequently, determination of catalytic parameters of ER Man I was not possible for site 4. The data, however, implied that *K*_*M*_ of ER Man I for site 4 is likely at least one order of magnitude higher than for all the other sites. Also for GnT I, site 4 showed the highest *K*_*M*_ value (table 1).

**Figure 3:**
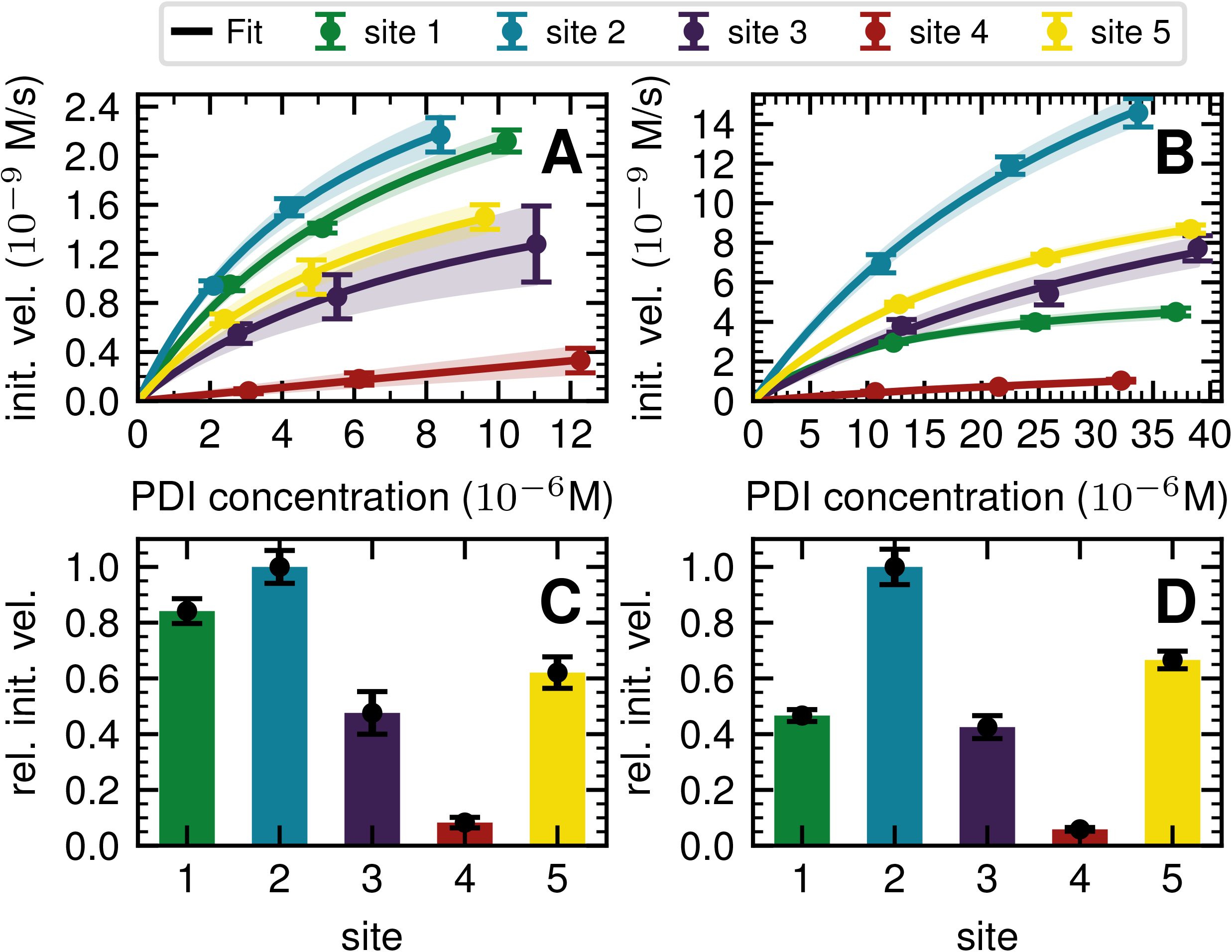
Michaelis-Menten analysis of processing kinetics. **A**: Initial velocities of ER Man I are plotted against three different substrate (PDI) concentrations and fitted to Michaelis-Menten kinetics by nonlinear regression (GraphPad Prism software). Error bars represent the standard error of linear regression fits in figure EV4 (*n*=1). The shaded error on the fit functions are individual fits to the upper and lower limit error range of the measured data points. **B:** Initial velocities of GnT I are plotted and fitted like described above for A. **C**: Relative initial velocities of ER Man I normalized by initial velocity of site 2. Initial velocities were averaged over three different PDI concentrations. Error bars are calculated by Gaussian error propagation of the shaded fit error in panels A and B. **D:** Relative initial velocities of GnT I normalized like described above for C.

**Table 1.** Apparent kinetic constants (± SEM) of ER Man I and GnT I for the five glycosylation sites of Man_9_GlcNAc_2_-PDI and Man_5_GlcNAc_2_-PDI, respectively.

|  |  | site 1 | site 2 | site 3 | site 4 | site 5 |
| --- | --- | --- | --- | --- | --- | --- |
| ER Man I | $V_{max}$ ($10^{-9}$ M/s) | $3.8 \pm 0.5$ | $3.7 \pm 0.3$ | $2.4 \pm 0.2$ | ND | $2.7 \pm 0.3$ |
| | $K_M$ ($10^{-6}$ M) | $8.1 \pm 1.8$ | $5.9 \pm 0.9$ | $9.4 \pm 1.3$ | ND | $7.5 \pm 1.3$ |
| | $k_{cat}$ ($s^{-1}$) | $6.3 \pm 0.8$ | $6.2 \pm 0.5$ | $3.9 \pm 0.3$ | ND | $4.4 \pm 0.4$ |
| | $k_{cat}/K_M$ ($10^6$ M<sup>-1</sup>s<sup>-1</sup>) | $0.8 \pm 0.2$ | $1.1 \pm 0.2$ | $0.4 \pm 0.1$ | ND | $0.6 \pm 0.1$ |
| GnT I | $V_{max}$ ($10^{-9}$ M/s) | $6.1 \pm 0.004$ | $31.4 \pm 4.1$ | $17.7 \pm 7.9$ | $3.2 \pm 1.3$ | $14.4 \pm 0.006$ |
| | $K_M$ ($10^{-6}$ M) | $13.2 \pm 0.02$ | $38.2 \pm 8.4$ | $52.2 \pm 37.1$ | $70.4 \pm 38.3$ | $25.1 \pm 0.02$ |
| | $k_{cat}$ ($s^{-1}$) | $0.1 \pm 0.0001$ | $0.5 \pm 0.1$ | $0.3 \pm 0.1$ | $0.05 \pm 0.02$ | $0.2 \pm 0.0001$ |
| | $k_{cat}/K_M$ ($10^3$ M<sup>-1</sup>s<sup>-1</sup>) | $6.8 \pm 0.1$ | $12.3 \pm 3.9$ | $5.0 \pm 4.5$ | $1.4 \pm 0.5$ | $8.0 \pm 0.01$ |

In addition to *K*_*M*_, *k*_*cat*_ values were calculated for ER Man I and GnT I, showing the highest turnover of substrate molecules on site 1 and 2, respectively. For both enzymes, *k*_*cat*_/*K*_*M*_ was highest for site 2, indicating that the glycan from site 2 was the preferred substrate of ER Man I and GnT I.

In order to compare initial velocities between sites, we normalized them by the initial velocity of site 2. These relative initial velocities were averaged over three PDI concentrations and reveal that the trimming of the oligosaccharide by ER Man I was approximately 10 times slower at site 4 compared to site 2 (figure 3C). For GnT I, site 4 reached only about 7% of the initial velocity of site 2 (figure 3D).

#### Rate constants describing appearance and disappearance of intermediate structures

Since GM I and GM II catalyze multiple, consecutive trimming reactions, each processing step was studied separately by recording the disappearance of the substrate, the transient accumulation of reaction intermediates and the formation of the final product. For fitting the dependence of the concentrations of all species on reaction time, we assigned apparent, first-order rate constants (k_1_-k_4_) to each reaction step as an approximation (figure 4A and B).

**Figure 4:**
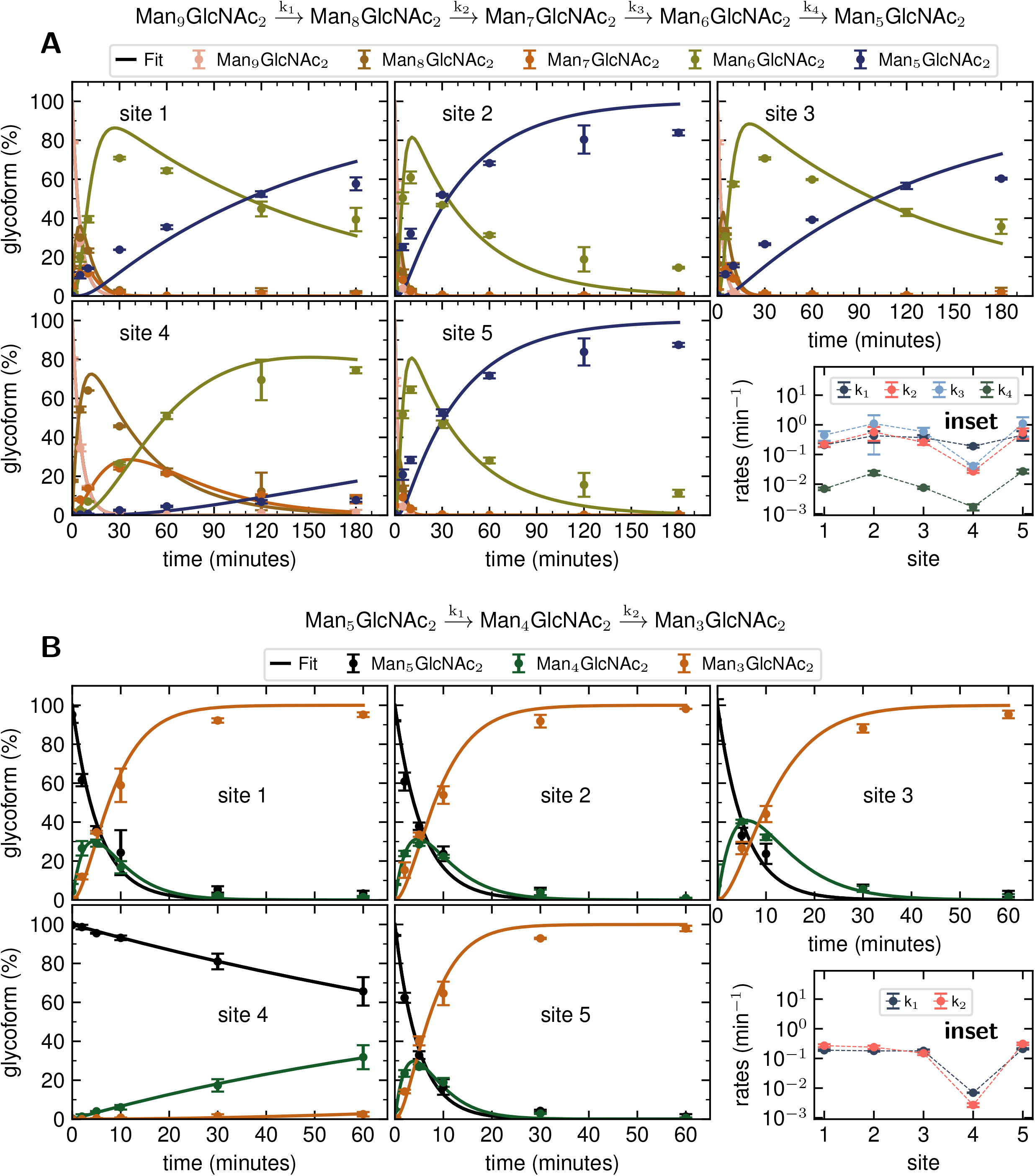
Site-specific processing by GM I and GM II. **A:** Consecutive mechanism for the step-wise removal of terminal mannoses from the Man_9_GlcNAc_2_ glycans of PDI by GM I. Conversion of Man_9_GlcNAc_2_ to Man_8_GlcNAc_2_, Man_8_GlcNAc_2_ to Man_7_GlcNAc_2_, Man_7_GlcNAc_2_ to Man_6_GlcNAc_2_, and Man_6_GlcNAc_2_ to Man_5_GlcNAc_2_ is described by apparent rate constants k_1_, k_2_, k_3_ and k_4_, respectively. Kinetics of glycan processing are shown for each of the five glycosylation sites of PDI individually. Solid lines represent a global fit of the data from each glycosylation site according to the mechanism shown above. Error bars represent the standard deviation of three independent experiments (*n*=3). **B:** Consecutive mechanism for the stepwise removal of terminal mannoses from the GlcNAcMan_5_GlcNAc_2_ glycans of PDI by GM II. Conversion of GlcNAcMan_5_GlcNAc_2_ to GlcNAcMan_4_GlcNAc_2_ and GlcNAcMan_4_GlcNAc_2_ to GlcNAcMan_3_GlcNAc_2_ is described by apparent rate constants k_1_ and k_2_, respectively.

For PDI (20 μM) glycan processing catalyzed by GM I (0.1 μM), the conversion of Man_9_GlcNAc_2_ to Man_8_GlcNAc_2_ (k_1_) occurred with similar kinetics at all sites. In contrast, values for k_2_ and k_3_ at site 4 were ~10−20 fold lower compared to those at the other sites (inset in figure 4A, table EV6), evidenced by the transient accumulation of the Man_8_GlcNAc_2_ and Man_7_GlcNAc_2_ intermediates (figure 4A). In addition, the results showed that the conversion of the Man_6_GlcNAc_2_ to the Man_5_GlcNAc_2_ glycoform was rate-limiting for the formation of the final product Man_5_GlcNAc_2_ at all glycosylation sites and occurred 16 to 34 fold slower than the slowest of the other processing steps. Again, site 4 showed the slowest Man_6_GlcNAc_2_ to Man_5_GlcNAc_2_ conversion with k_4_ being 4-16 fold lower than the k_4_ values at the other sites.

The global fits of Man_9_GlcNAc_2_ processing at sites 1-3 and 5 by GM I agreed reasonably well with the experimental data (figure 4A). The largest deviations from the fit were observed for the kinetics of formation of the final product Man_5_GlcNAc_2_. This indicated that the processing mechanism might be more complex than a consecutive 4-step mechanism and might include branch points and parallel pathways [24].

We therefore extended the reaction mechanism by adding a branch point after Man_7_GlcNAc_2_, assuming the formation of two different Man_6_GlcNAc_2_ isomers (appendix figure S1, appendix table S1). The corresponding six-parameter fit indeed agreed better with the experimental data, but we consider the results underdetermined, because we could not experimentally distinguish the different Man_6_GlcNAc_2_ isomers.

As shown above for GM II (figure 2D), hardly any conversion of GlcNAcMan_5_GlcNAc_2_ to GlcNAcMan_3_GlcNAc_2_ could be observed on site 4. To determine the affected mannose trimming step, we measured kinetics of GM II (67 nM) for the processing of the substrate GlcNAcMan_5_GlcNAc_2_-PDI (20 μM). The data of each glycosylation site was fitted according to a consecutive mechanism with two apparent, first-order rate constants k_1_ and k_2_ approximating the conversion of GlcNAcMan_5_GlcNAc_2_ to the intermediate GlcNAcMan_4_GlcNAc_2_ and the product GlcNAcMan_3_GlcNAc_2_, respectively (figure 4B). At sites 1-3 and 5, both trimming steps occurred with practically identical rates (inset in figure 4B, table EV7), the substrate was consumed within 30 minutes and the intermediate reached a maximum level of ~30% after five minutes. In contrast, both steps were significantly slower at site 4, with k_1_ and k_2_ being ~30 fold and 50-100 fold lower compared to the other sites. Consequently, the intermediate had only gradually accumulated to ~30% after 60 minutes of the reaction (figure 4B).

#### Influence of the protein structure on site-specific mannose trimming

Our atomistic MD simulations revealed an essential role of the tertiary structure context for the site-specific substrate properties of identically composed glycans. PDI contains two catalytic disulfide bonds and one additional, structural disulfide bond (figure 1B) [30, 34]. Nearly complete formation of these disulfide bonds (96%) was confirmed by Ellman assay. To probe the influence of glycoprotein structure on site-specific processing kinetics, we altered the structure of PDI by reducing and subsequently alkylating the three disulfide bonds. The alkylation resulted in slight conformational changes as detected by far-UV circular dichroism (CD) spectroscopy (figure 5A). As this structural change was by no means comparable to that occurring upon complete denaturation of PDI with 6 M guanidine hydrochloride, the structure of reduced and alkylated PDI still remained native-like.

**Figure 5:**
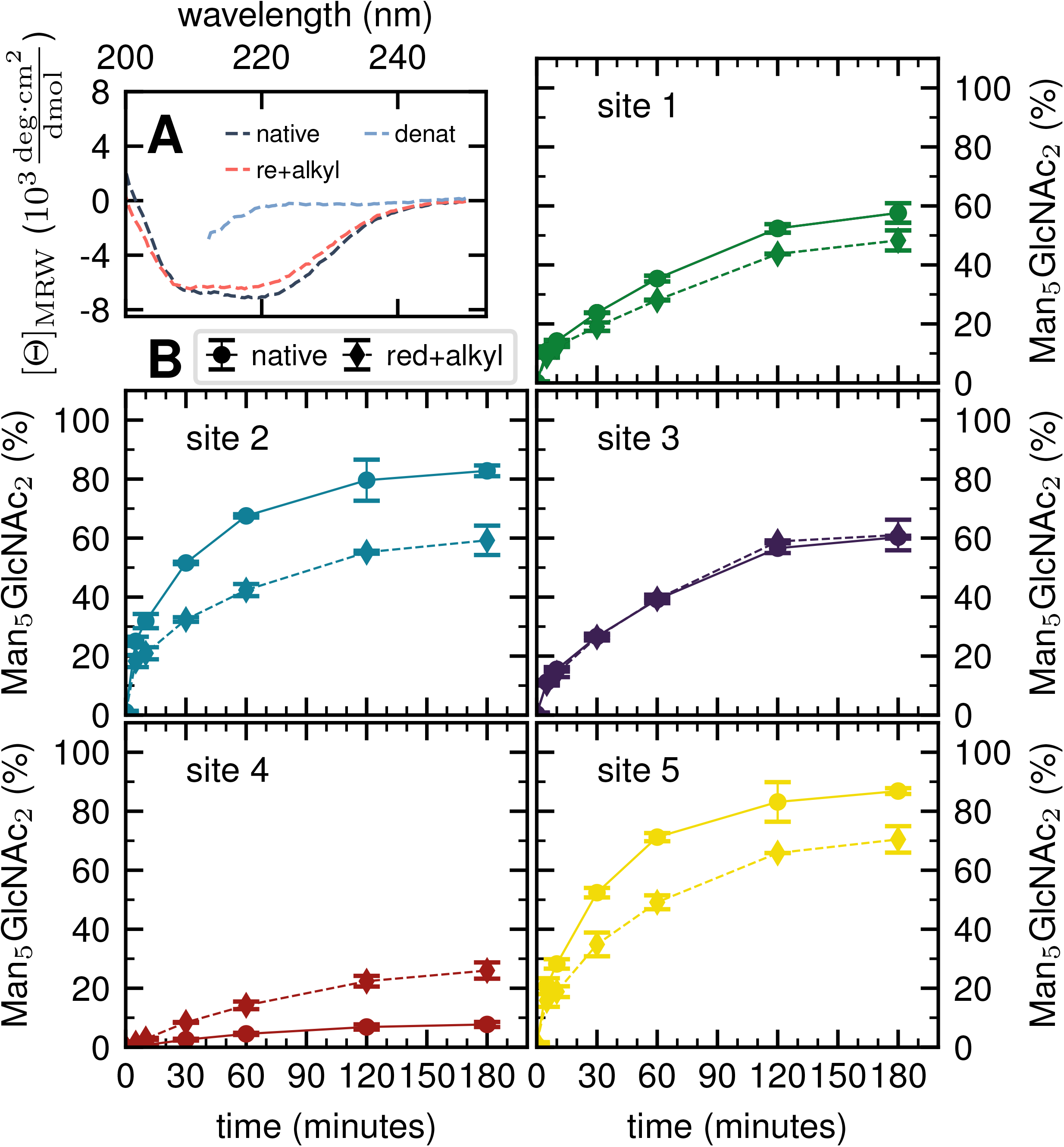
Influence of protein structure on site-specific glycan processing by GM I. **A:** Far-UV CD spectra of native, reduced and alkylated, and denatured PDI. Reduction and alkylation of the three disulfide bonds led to a slight change in the structure of PDI. The less pronounced minimum (at 222 nm) and maximum (at 200 nm) indicated a small loss of secondary structure. **B:** Reduced and alkylated as well as native PDI was incubated with GM I. Shown is the conversion of Man GlcNAc to Man GlcNAc on each site over 180 minutes in comparison to processing of the respective site on native PDI. Error bars represent the standard deviation of three independent experiments (*n*=3).

Reduced and alkylated as well as native, oxidized Man_9_GlcNAc_2_-PDI (20 μM) was then used as a substrate for GM I (0.1 μM) (figure 5B). Site-specific formation of the final product Man_5_GlcNAc_2_ was monitored and compared to the respective site on native PDI. We observed that site 4 was processed more efficiently on the reduced and alkylated protein, whereas site 1, 2 and 5 glycans were hydrolyzed slower. For the site 3 glycan, no difference in processing by GM I was observed.

## Discussion

We demonstrated that for a given enzyme, glycan processing at the five sites of PDI occurred with different kinetics. For all the enzymes tested, site 4 was processed the slowest, while sites 2 and 5 were processed most efficiently. This order of reactivity was also observed for the same glycoprotein *in vivo* in insect and CHO cells [15, 19]. We therefore concluded that identical glycans at different glycosylation sites presented different substrate properties to the processing enzymes.

We used MD simulations to study the glycan conformations and accessibility on the protein surface in order to explain the site-specific substrate properties of the *N*-linked glycans. An effective enzyme accessibility of ‘free’ and/or ‘exposed’ glycan conformations exhibited a clear site-specificity of the substrate availability and can be qualitatively compared to the enzyme accessibility through the particular structure and glycan-enzyme binding modes. For instance, GH47 α-mannosidases typically have a deep and narrow funnel to the catalytic center that binds one glycan branch at a time [33]. Hence, the contacting conformations, which expose only individual branches, may still be accessible for enzyme processing. The convex and concave protein surface topologies in the immediate surrounding of site 2 and 4 (figure EV 1) represent different presentation platforms of the *N*-glycan that also affect the glycan accessibility. At site 2, the positive protein surface curvature potentially minimizes steric clashes when binding to α-mannosidase. In contrast, the concave protein surface around site 4 potentially interferes with binding by the enzyme even though the glycan conformation might not be in contact with the protein surface. Stretched and extended conformations of glycans may be ideally suited for binding in the deep binding pocket of α-mannosidases. However, at site 4 the steric hindrance of a concave protein surface diminishes the accessibility of the otherwise ‘free’ glycan conformations for processing enzymes. At site 2, the combination of a convex protein surface topology and extended glycan branches may be especially advantageous for unhindered glycan-enzyme complexation. This preference of site 2 over site 4 was observed experimentally and we therefore concluded that the substrate properties of *N*-linked glycans as observed by MD simulations reflected their biochemical properties.

An *N*-linked glycan on a convex protein surface represents an ideal substrate for a given enzyme, with the affinity solely determined by the glycan-enzyme interaction. Indeed, for ER Man I and GnT I we found the site 2 glycan to be the preferred substrate (highest *k*_*cat*_/*K*_*M*_ value). However, the relative initial reaction rates for the other sites differed between the two enzymes, showing that the interpretation of the ‘in contact & exposed’ glycan conformations in terms of accessibility as well as the influence of the protein surface topology depends on the exact glycoprotein-enzyme complex. For site 4, we noted a higher *K*_*M*_ value for both enzymes, indicative for additional protein-glycan interactions due to the concave nature of the protein surface at this glycosylation site. Accordingly, the structural differences of ER Man I and GnT I explain the different site-specific effects on the respective *K*_*M*_ values. When the atomistic details of these complexes will hopefully become available in the future, we anticipate that a quantitative and qualitative analysis of glycans based on MD simulations can be further refined.

We perturbed glycan-protein interactions by alkylation of PDI. Similarly, denaturing the glycoprotein soybean agglutinin with 8 M Urea improved the processing of its *N*-glycans by ER Man I [35]. However, our data on reduced and alkylated PDI suggest that subtle structural differences can already have site-specific effects: well processed and therefore probably easily accessible *N*-glycans from sites 1, 2 and 5 showed slower processing kinetics upon alkylation. We hypothesize that the change in protein structure featured new interactions, not present in the native PDI, between the glycans from sites 1, 2 and 5 and the PDI surface. For the site 4 glycan, on the other hand the slight change in protein structure improved processing significantly, while there was no effect detectable for site 3.

Within the framework of our hypothesis, folding intermediates in the ER or conformational isomers of folded proteins in the ER and Golgi may represent distinct substrates for site-specific glycan processing in kinetically controlled processing pathways. The differential processability of a defined *N*-linked glycan might even display the folding status of the covalently linked protein in processes such as the quality control pathway of protein folding in the ER [5].

Our detailed biochemical analysis allowed us to follow enzymes that perform multiple processing steps like GM I and GM II. We identified the rate-limiting step in trimming of a Man_9_GlcNAc_2_ to Man_5_GlcNAc_2_ by GM I to be the last step (Man_6_GlcNAc_2_ to Man_5_GlcNAc_2_). Lal *et al.* argued that the last mannose trimmed by GM I is the terminal mannose from the B-branch of the glycan, which is *in vivo* trimmed in the ER by ER Man I [24]. In case of PDI, this last trimming step was greatly impaired on site 4.

Trimming of two mannoses from GlcNAcMan_5_GlcNAc_2_ by GM II is an essential step in the conversion of hybrid to complex *N*-glycans [36]. Our analysis showed that GM II has, compared to the other enzymes tested, the lowest activity on site 4 with a decrease in activity of two orders of magnitude relative to the other sites, which explained why a secreted version of PDI produced in CHO cells showed the highest percentage of hybrid glycan structures on site 4 [19]. However, our analysis showed that GM II was able to process the site 4 glycan to some degree, indicating a time and/or enzyme-limited process *in vivo.*

The eukaryotic secretory pathway is organized such that glycoproteins are exposed for a limited time to processing enzymes located in different compartments of the pathway [37–39]. Therefore, site-specific initial processing velocities are a determining factor for *N*-glycan processing *in vivo*, together with the residence times of glycoproteins in the individual compartments of the secretory pathway. In such a kinetically controlled system, small alterations of enzyme levels, as observed for example during cellular differentiation in a multicellular organism, can have strong qualitative and quantitative effects on the glycoproteome. Site-specific glycan structures will be affected differently on proteins with multiple glycosylation sites. Therefore, a quantitative glycoproteomics analysis will become a necessity in order to understand the functional properties of glycoproteins.

## Material and methods

### MD simulations

The crystal structure from the Protein Data Bank (PDB: 2B5E [30]) was used as template for PDI. The simulations with the full-length PDI glycoprotein were prepared to account for the flexible nature of the glycans and the protein. The glycoprotein conformations were pre-sampled with the REST2 Hamiltonian replica exchange method [40] as described in Section S.1. The enhanced conformational sampling was used in two ways: (i) construction of an optimized triclinic box size [41] as described in Section S.2, and (ii) the sampled structures served as starting configurations for the production runs.

All MD simulations were performed using the GROMACS 2018 simulation software [42]. The AMBER ff99SB-ILDN force field [43] was used for the protein and the GLYCAM06i force field [44] for the glycans. One setup contained the full-length PDI protein with Man_9_GlcNAc_2_ glycans attached at Asn82 (site 1), Asn117 (site 2), Asn151 (site 3), and Asn170 (site 4). This setup was solvated in 63’000 TIP5P water molecules [45] in the optimized box (see Section S.2), and 42 sodium ions [46] were added to neutralize the system. A second setup only contained the a’-domain with a Man_9_GlcNAc_2_ glycan at Asn425 (site 5), starting from residue Lys366, which was capped by a N-Me amide group. This setup was solvated by 12’000 TIP5P water molecules in a 6.4 nm x 6.4 nm x 9.5 nm rectangular box such that the glycan was pointing along the z-direction. For system neutralization, this setup contained 11 sodium ions [46].

All MD simulations were performed using the leapfrog integrator [47] with a step size of 2 fs. The solute coordinates were stored every 20 ps. The enhanced sampling and production protocols were carried out under an isothermal-isobaric (NPT) ensemble with periodic boundary conditions. The temperature was kept constant at T = 300 K using the V-rescale thermostat [48] with a coupling time of 0.1 ps. The pressure was kept constant at 1 bar using the Parrinello-Rahman barostat [49] with a coupling time of 2 ps and a compressibility of 4.5 × 10^−5^bar^−1^. The Verlet cutoff-scheme [50] was applied for non-bonded interactions with a cutoff of 1.0 nm, a neighbor list update frequency of 20 fs, and a buffer tolerance of 10^−4^. The smooth particle mesh Ewald method [51] with a grid spacing of 0.16 nm and interpolation order of four was used for long-range electrostatics. The SHAKE algorithm [52] was applied to rigidify all bonds.

For the exhaustive sampling of the glycoprotein ensemble, random frames were chosen from the REST2 replica trajectory at T = 300 K. The glycans in these conformations were removed and new Man_9_GlcNAc_2_ glycans were attached using the doGlycans tool [53]. These glycan starting conformations were randomly varied within the thermodynamically most stable configurations. The thermodynamically most stable glycosidic bond dihedral angles were taken from Ref. [54] and randomly varied within a range ±40° upon a particular MD starting setup. The generated glycoprotein conformations were solvated in the previously optimized box. The first 10 ns of the simulations were discarded as equilibration. The final production trajectories ranged from 100 ns to 400 ns.

### Markov state modeling

Inverse distances of glycan residues to vicinal hexoses and amino acids (with hydrogen bond donating/accepting side chains) were chosen as features for the MSMs, as described before [13, 14]. Each glycan was individually analyzed using a site-specific choice of considered neighboring glycans and protein surface residues for the (inverse) distance calculations (table EV1). For the distances involving hexoses, the respective O5 atom positions were taken. For amino acids, a side-chain specific atom position was considered (listed in table EV2). Note that in contrast to the previous MSMs of glycoproteins [13, 14], where very few features were manually selected for the sake of modeling ease, our featurization includes all spatially reachable amino acid and carbohydrate moieties.

In the next step, a dimensionality reduction was performed using the principal component analysis (PCA) and choosing the first ten dimensions (n = 10). Subsequently, the hierarchical volume-scaled common nearest neighbor (vs-CNN) clustering algorithm was used to discretize the trajectory into conformational microstates, i.e. clusters [55]. We started by finding an initially large cutoff R_0_, such that 99% of the data was clustered, while the similarity N = 10 was fixed. For sites 1-5 different values were respectively obtained for R_0_ = {1.4, 1.2, 1.4, 1.4, 1.0}. Next, the cutoff was decreased in steps of R_i+1_ = R_i_ · e^−β·Δ F/n^ where ΔF is the free energy difference between hierarchical levels and β = 1/k_B_T with k_B_ as Boltzmann constant. Several hierarchical trees were tested using the parameter values ΔF = {0.6, 0.8, 1.0, 1.2, 1.5, 2.0} k_B_T. In each step *i*, a given cluster of minimum size *N*_split_ was updated if it was split into at least two new clusters of minimum size *N*_keep_. These parameters were chosen as pairs from N_split_ = {101, 1001} and N_keep_ = {11, 101, 1001}, while N_split_ ≥ N_keep_ was ensured. The transition matrix of a particular microstate state-space was estimated using milestoning in the maximum likelihood estimation from the PyEMMA package [32]. Since density-based clustered data was previously shown to improve the Markovian assumption for core-set MSM building [56–58], a lag time of 100ps could be chosen in this work. The major quantity obtained from the MSM is the stationary probability P_i_ of microstate *i* contained in the first eigenvector of the transition probability matrix [56–58].

### Accessibility assessment

For each microstate the glycan conformations were quantified by the solvent accessible surface area (SASA) A_solv_, the number of hydrogen bonds N_h_, and the number of atomic contacts N_c_ with the site-specific glycoprotein environment. The SASA was determined using the Shrake-Rupley algorithm implementation in MDTraj [59, 60]. The probe sphere radius was set to 1.4 Å and 960 points were used to represent the surface of an atom. For the hydrogen-bond count N_h_, a geometric criterion of 0.35 nm donor-acceptor distance and a hydrogen-donor-acceptor angle of 30° was used. For consistency, the contact criterion for N_c_ also involved an atomic distance threshold of 0.35 nm. During the hydrogen bond and contact analysis of a given mannose residue, the direct neighboring mannoses were excluded. Also, the hydrogen bond count was subtracted from the number of atomic contacts for each frame to avoid double counting.

The above quantities were averaged across the conformations in a microstate *i* for each mannose *k*, *l*, and *m* of branch *Y*, such as, *k* = 9, *l* = 6, and *m* = 4 of branch A (see figure 1 A). The average of the full length of branch *Y* was the sum ⟨·⟩_Y,i_ = ⟨·⟩_k,i_ + ⟨·⟩_*l*,*i*_ + ⟨·⟩_*m*,*i*_. Generally, varying lengths of the branches could be compared whilst taking one {*k*}, two {*k*, *l*}, or three {*k*, *l*, *m*} monosaccharides into account. In the remainder, we use the subscripts in the notation for the average ⟨·⟩_Y,i_ only if required.

To quantify the accessibility or exposure of a given monosaccharide in a glycan branch, the exposure score *s*_x_ (x-score) was defined as the combination of the individual measures as follows,

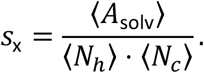

where ⟨*A*_solv_⟩ is the average SASA value, ⟨*N*_ℎ_⟩ the average number of hydrogen bonds, and ⟨*N*_*c*_⟩ the average number of contacts. The x-score was calculated for each glycan branch separately, considering the varying length of the branches.

We classified ‘free’ and ‘contact’ microstates via the number of atomic contacts across all full-length glycan branches, i.e., 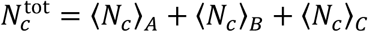. For a given threshold 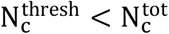 the glycan conformations within the microstate were considered in ‘contact’. Hence, for 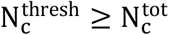 the glycan conformations of the microstate were considered to be ‘free’. Additionally, branch *Y* with x-score 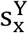 in a given microstate was considered ‘exposed’ if for a given threshold 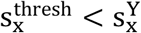. In combination with the MSMs, the relative abundance of particularly classified conformations was calculated as the ensemble average based on the stationary probability distribution from the MSMs. For example, the fraction of exposed conformations is 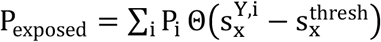 where Θ(x) is the Heaviside step function. Also, combinations of ‘free’ and ‘contact’ with the ‘exposed’ classifications, respectively, were considered, as 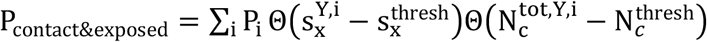. These ensemble averages and their standard deviation of microstate classification were calculated over several MSMs, i.e., stationary probability distributions P_i_ from the different clustering results (section Markov state modeling), and varying threshold values of 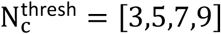 and 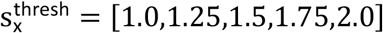.

### Constructs

From the PDI expression construct pRG85 [15], the C-terminal sequence “LELQLEHDEL” was removed by using the primers prHI_14_fw and prHI_15_rev (table appendix S2). PCR was performed with Phusion polymerase (Thermo Fisher) and phosphorylated primers. PCR products were ligated with T4 ligase (NEB). The resulting plasmid (pRG85_CM) was transformed into DH5α *E. coli* cells. Baculovirus stocks for secreted, *N*-terminally His_8_-tagged, *human* glycan processing enzymes ER Man I (MAN1B1), GM I (MAN1A2) and GnT I (MGAT1) were obtained from the glycoenzyme repository http://glycoenzymes.ccrc.uga.edu/ [61]. Purified, recombinantly produced *Drosophila melanogaster* Golgi Mannosidase II (α-Man-II a) was kindly provided by Dr. Douglas Kuntz from the University of Toronto [62].

### Production and purification of substrate PDI

Recombinant bacmids containing the gene of interest (PDI) were generated according to the Baculovirus Expression system manual (Thermo Fisher Scientific, Cat. No. 10359016). In brief, DH10Bac *E. coli* cells (Thermo Fisher, Cat. No. 10361-012) were transformed with pRG85_CM and bacmid DNA was subsequently isolated and used to transfect Sf21 cells (Thermo Fisher, Cat. No. 11497013) using the Cellfectin™ II reagent (Thermo Fisher, Cat. No. 10362100). Recombinant viruses were harvested after 72 hours from the supernatant. Expression and purification of PDI was done as described by Hang *et al.* [15]. Briefly, High-Five™ cells (Thermo Fisher, Cat. No. B855-02) were infected with the virus stock. For homogenously glycosylated PDI (Man_9_GlcNAc_2_ on all sites) 10 μM α-1, 2-mannosidase inhibitor kifunensine (Sigma-Aldrich Chemie GmbH, Cat. No. K1140) was added to the cell culture. After 48 hours, cells were lysed and the His_10_-tagged PDI was purified using Ni-NTA beads (Protino, Cat. No. 745400.100). Purified PDI was buffer exchanged to the activity buffer required by the enzyme used in the assay. PDI concentration was determined using the NanoDrop UV-Vis spectrophotometer (Thermo Fisher). For storage, PDI was flash-frozen in liquid nitrogen and kept at −80°C. Figure appendix S2 shows that freezing and thawing of PDI had no influence on the site-specific processing of its *N*-glycans.

### Production and purification of glycan processing enzymes

High-Five™ cells were infected with recombinant baculovirus stocks for either ER Man I, GM I or GnT I like described before for PDI. After 72 hours, the cell culture was centrifuged at 3500 rcf for ten minutes, supernatants were filtered using 0.22 μM filters (TRP, Cat. No. 99722) and enzymes were purified *via* their His-tag. Therefore, 1 ml of Ni-NTA beads per 50 ml supernatant were washed three times with 10 column volumes (CV) phosphate buffered saline (PBS) and subsequently mixed with the supernatant. Batch binding was done on a spinning wheel for three hours at 4°C. The bound fraction was transferred to a gravity flow column (Macherey-Nagel, Cat. No. 745250.10) and washed with 15 CV washing buffer (25 mM Imidazole in PBS, adjusted with HCI to pH 7). Elution was done with 4 CV of elution buffer (250 mM imidazole in PBS, pH 7). Purified enzymes (figure appendix S3) were buffer exchanged to PBS with 10% Glycerol (v/v), flash frozen in liquid nitrogen and stored at −80 °C.

### Enzyme *in vitro* assays

All *in vitro* assays were performed with PDI as a substrate and either ER Man I, GM I, GnT I or GM II in an enzyme specific activity buffer (table appendix S3). Immediately after addition of the enzyme to PDI, the reaction mix was incubated at 37°C and shaken at 500 rpm.

For *in vitro* assays with ER Man I and GM I, PDI purified from High-Five™ cells treated with kifunensine was used as a substrate. PDI therefore showed mainly Man_9_GlcNAc_2_ on all sites, presenting a homogenous glycosubstrate for the tested enzyme (figure EV3A).

For the GnT I *in vitro* assay purified PDI (without kifunensine treatment) was pre-incubated over night with the enzyme GM I (enzyme to substrate molar ratio 1:200) at room temperature to hydrolyze glycans (Man_9_GlcNAc_2_ to Man_6_GlcNAc_2_) to Man_5_GlcNAc_2_, the glyco-substrate of GnT I (figure EV3B). The GnT I assay was subsequently performed in GnT I activity buffer [63] containing 5 mM of UDP-GlcNAc (Sigma-Aldrich Chemie GmbH).

The substrate preparation for the GM II assay was done by successively incubating PDI (without kifunensine treatment) with two glycan processing enzymes. First with GM I (enzyme to substrate molar ratio of 1:300) for four hours at 37°C to obtain mainly Man_5_GlcNAc_2_ on all sites. After buffer exchange to GnT I activity buffer, 5 mM of UDP-GlcNAc and purified GnT I in an enzyme to substrate molar ratio of 1:300 was added to the reaction mix. After four hours of incubation at 37°C PDI showed mainly GlcNAcMan_5_GlcNAc_2_ on all sites (figure EV3C).

At indicated time points, samples containing a minimum of 50 μg of PDI were taken. To stop the reaction each aliquot was mixed with trichloroacetic acid (15% final concentration) and kept on ice for five minutes. PDI was subsequently pelleted, washed and stored as described by Hang *et al.* [15].

### Sample preparation and MS measurement

Protein pellets were dissolved in 50 μl of 8 M Urea and loaded onto a 30 K centrifugal filter unit (Merck). Preparation for MS was done as described in [64]. Briefly, the protein was reduced by 50 mM dithiothreitol and alkylated by 65 mM iodoacetic acid in 0.05 M ammonium bicarbonate buffer (pH 8.5) to facilitate Trypsin digestion (Trypsin to PDI weight ratio 1:50) at 37°C for 16 h. For analysis of glycosite occupancy, Trypsin digest of PDI was followed by digestion of *N*-glycans with endo-β-*N*-acetylglucosaminidase H (500 U; NEB) in sodium citrate buffer (50 mm, pH 5.5) at 37 °C for 40 hours [65]. Peptides were collected by centrifugation and desalted by Zip-Tip C18. For analysis, samples were dissolved in 2% acetonitrile with 0.1% formic acid and analyzed by one of the two methods described below:

Either by a calibrated LTQ-Orbitrap Velos mass spectrometer (Thermo Fischer) coupled to an Eksigent-Nano-HPLC system (Eksigent Technologies) like described in [15].

Alternatively, by a calibrated Q Exactive™ mass spectrometer (Thermo Fischer) coupled to a Waters Acquity UPLC M-Class system (Waters) with a Picoview™ nanospray source 500 model (New Objective). Samples were loaded onto a Acclaim PepMap 100 trap column (75 μm × 20 mm, 100 Å, 3 μm particle size) and separated on a nanoACQUITY UPLC BEH130 C18 column (75 μm × 150 mm, 130 Å, 1.7 μm particle size), at a constant flow rate of 300 nL/min, with a column temperature of 50 °C and a linear gradient of 1−35% acetonitrile/0.1% formic acid in 42 min, followed by a sharp increase to 98% acetonitrile in 2 min and then held isocratically for another 10 min. One scan cycle comprised of a full scan MS survey spectrum, followed by up to 12 sequential HCD scans based on the intensity. For glycosylation profiling analysis, full-scan MS spectra (800–2000 m/z) were acquired in the FT-Orbitrap at a resolution of 70,000 at 400 m/z, while HCD MS/MS spectra were recorded in the FT-Orbitrap at a resolution of 35,000 at 400 m/z. HCD MS/MS spectra were performed with a target value of 5e^5^ by the collision energy setup at a normalized collision energy 22.

### Glycoform quantification

Spectra obtained were analyzed with XCalibur 2.2 sp1.48 (Thermo Fisher) like described previously [15]. For quantification, extracted ion chromatography of all glycoforms were plotted by their unique mass over charge (m/z) ratio, based on a previous study by Losfeld *et al.* (table appendix S4) [19]. Peak area was defined manually and integrated by the program. The relative amount of each glycoform located on the same peptide backbone was calculated as shown before by Hang *et al* [15].

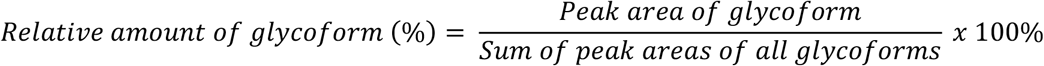

### Kinetics of site-specific glycan processing by GM I and GM II

For GM I, the glycan processing kinetics of each site were globally fitted using Dynafit [66] according to the following consecutive mechanism

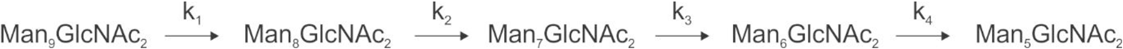

where the apparent rate constants k_1_, k_2_, k_3_ and k_4_ describe the consecutive trimming of one terminal mannose residue from the PDI-Man_9_GlcNAc_2_ substrate. For GM II, the glycan processing kinetics of each site were globally fitted using OriginPro 2018b (OriginLab) according to the 2-step consecutive mechanism

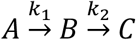

where A, B and C represent PDI-GlcNAcMan_5_GlcNAc_2_, PDI-GlcNAcMan_4_GlcNAc_2_ and PDI-GlcNAcMan_3_GlcNAc_2_. The fractions of these glycoforms are then given by

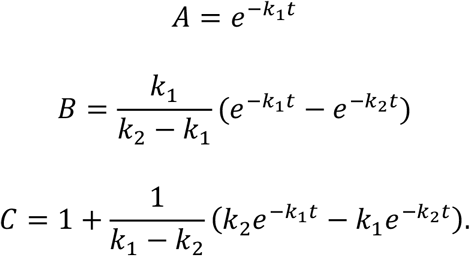

The apparent rate constants k_1_ and k_2_ were shared among the three datasets of each glycosylation site.

### Reduction and alkylation of PDI

Reduction and alkylation of PDI was done as described earlier for preparation of proteins for MS [64]. The alkylation of cysteines was highly efficient as only alkylated peptides were detected by MS.

### Circular dichroism

To determine if reduction and alkylation of the protein changed the protein structure, far-UV circular dichroism (CD) spectra were recorded. PDI was purified from kifunensine treated High-Five™ cells. To increase purity of the PDI sample two additional purification steps on a Basic Äkta module (Amersham Biosciences) were introduced. First, PDI (in PBS, pH 7) was submitted to size exclusion chromatography on a Superose™ 12 10/300 GL column (GE Healthcare, Cat. No. 17-5173-01) at a flow rate of 0.5 ml/min with PDI eluting between 11 and 14 ml.

In the second step, hydrophobic interaction chromatography using a Hi-Trap Butyl-Sepharose column (GE Healthcare, Cat. No. 28411001) with a flow rate of 1 ml/min and a gradient from 0.9 M (NH_4_)_2_SO_4_ in PBS to PBS over 5 CV was performed. The peak eluting between 16 and 22.5 ml was collected, and buffer exchanged to PBS.

Far-UV CD spectra of oxidized native, reduced and alkylated and denatured oxidized PDI (with 6 M guanidine hydrochloride) were recorded at a protein concentration of 0.4 mg/ml in PBS, pH 7.0 at 25 °C using a temperature-controlled J715 CD spectrometer (Jasco). CD signals were converted to mean residue ellipticity as described by Schmid *et al.* [67].

## Acknowledgment

We thank Professor Donald Jarvis and Professor Kelley Moremen who developed the “Glycoenzyme repository” from which we obtained the expression constructs for the enzymes ER Man I, GM I and GnT I. Additionally, we would like to thank Dr. Douglas Kuntz for kindly providing us with purified GM II enzyme. We also thank the Functional Genomics Centre Zürich for their help with the MS analysis. We thank Tsjerk Wassenaar for providing the source code for the near-densest lattice packing algorithm and the corresponding rotational motion removal in GROMACS 2018. This project was supported by the Swiss National Supercomputing Centre (CSCS) under Project ID s895, the Swiss National Science Foundation Grants 310030_162636 and 310030B_182835 to MA and ETH Zurich.

## Author contributions

MA and SR designed the research. MEL, CM, and RGW designed the experiments and simulations. CM and CG performed the experiments. RGW performed the simulations. CM, RGW, CG, RG and CWL analyzed the data. CM, RGW, CG, RG, SR, and MA wrote the manuscript. All authors read and approved the final manuscript.

## Expanded View Figure Legends

**Figure EV 1:**
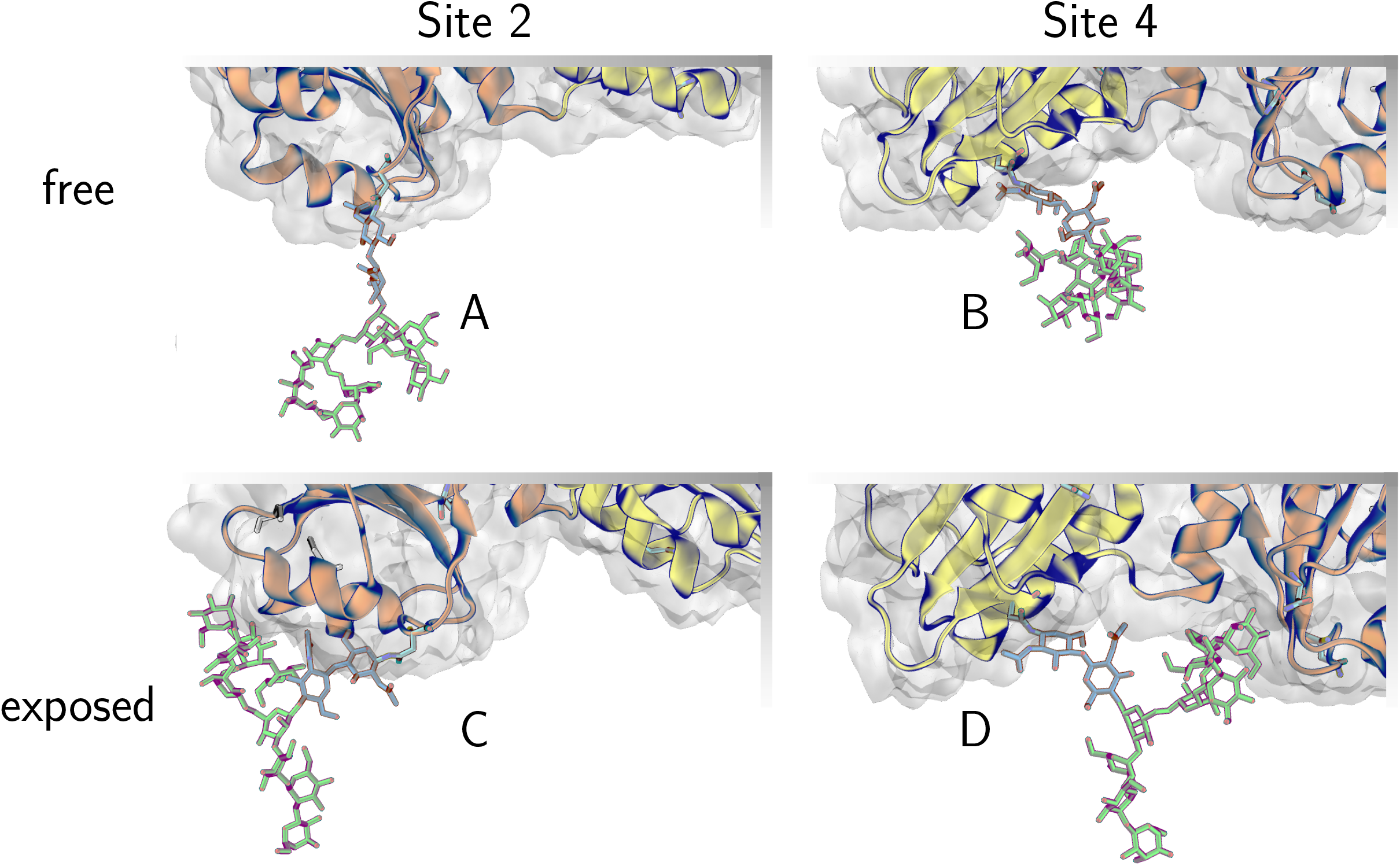
Convex and concave protein surface on sites 2 and 4. The convex and concave protein surface at sites 2 and 4 represent different glycan accessibility. **A:** The positive curvature of the protein surface at site 2 allows extended branch conformations if the whole glycan is classified as ‘free’. **B:** In contrast, the glycan at site 4 is confined by the concave protein surface environment, such that even ‘free’ conformations are still dominated by collapsed conformations. Hence, at site 4 the steric hindrance of a concave protein surface diminishes the accessible space for ‘free & exposed’ glycan conformations **C:** Extended branch conformations are particularly ‘exposed’ at site 2 even if the glycan is in ‘contact’. **D:** While at site 4 individually ‘exposed’ branches are still relatively buried inside the concave protein environment if the glycan is in ‘contact’ with the protein surface.

**Figure EV 2:**
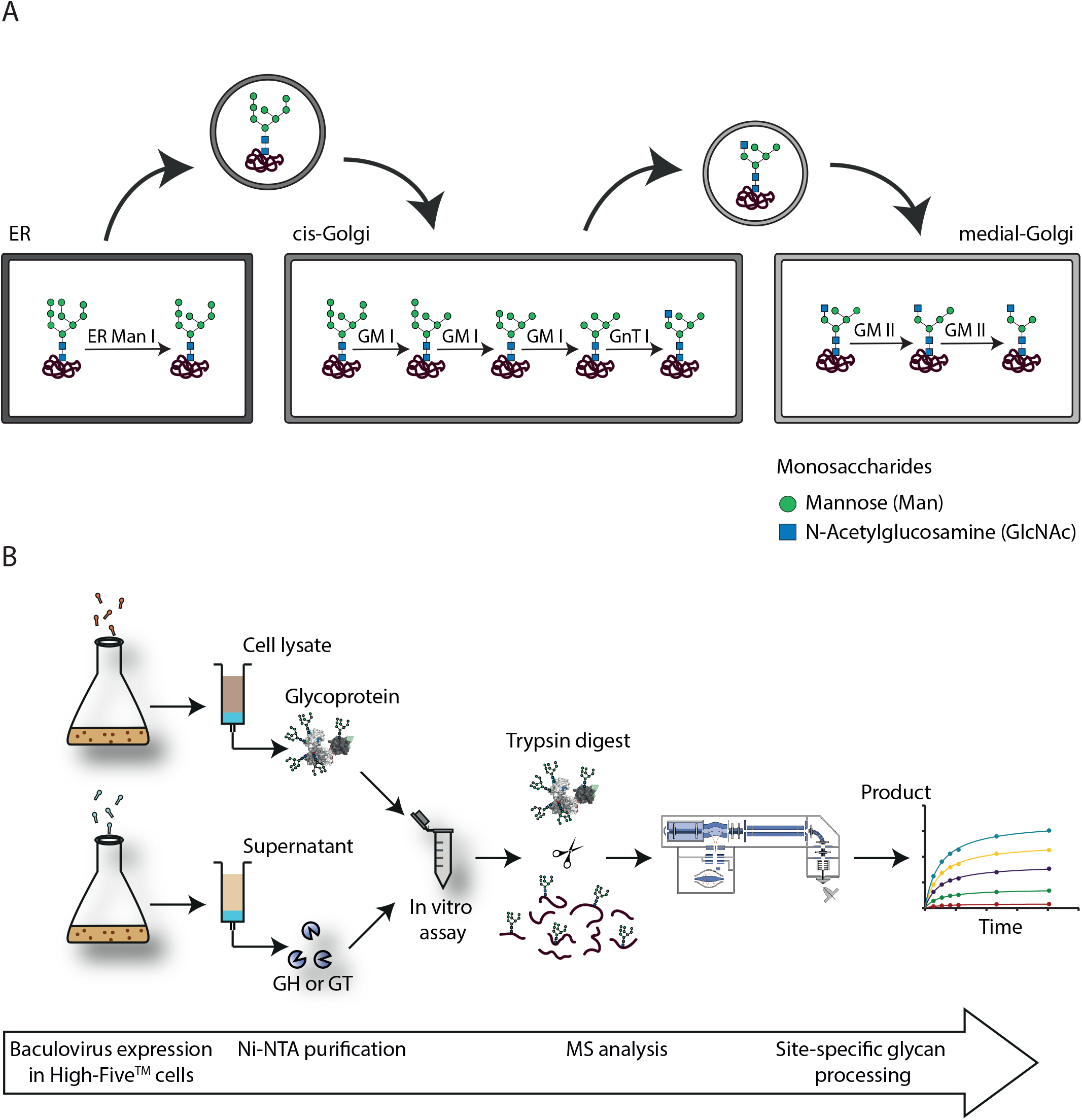
*N*-glycan processing *in vivo* and *vitro*. **A:** Simplified representation of the *N*-glycan processing steps in mammalian cells. Shown are only the reactions relevant to this study, catalyzed by ER Man I, GM I, GnT I and GM II. Vesicles transport the processed glycoproteins between the different compartments (ER, cis-Golgi and medial-Golgi). This transport defines the time glycoproteins are exposed to the respective glycan-processing enzymes [8]. **B:** Workflow of *in vitro* assays to study site-directed glycan processing. The model protein PDI and different glycan processing enzymes, glycosylhydrolases (GH) and a glycosyltransferase (GT), were produced in High-Five™ cells using the Baculovirus expression system. PDI was retained in the ER of the cells and therefore purified from the cell lysate. Glycan-processing enzymes were secreted and therefore purified from the supernatant with metal chelate (Ni-NTA) affinity chromatography. For the *in vitro* assay PDI and the respective enzymes were incubated, and the reactions were stopped by TCA precipitation after different incubation times. Tryptic glycopeptides were analyzed by MS and product formation over time was quantified individually for each site.

**Figure EV 3:**
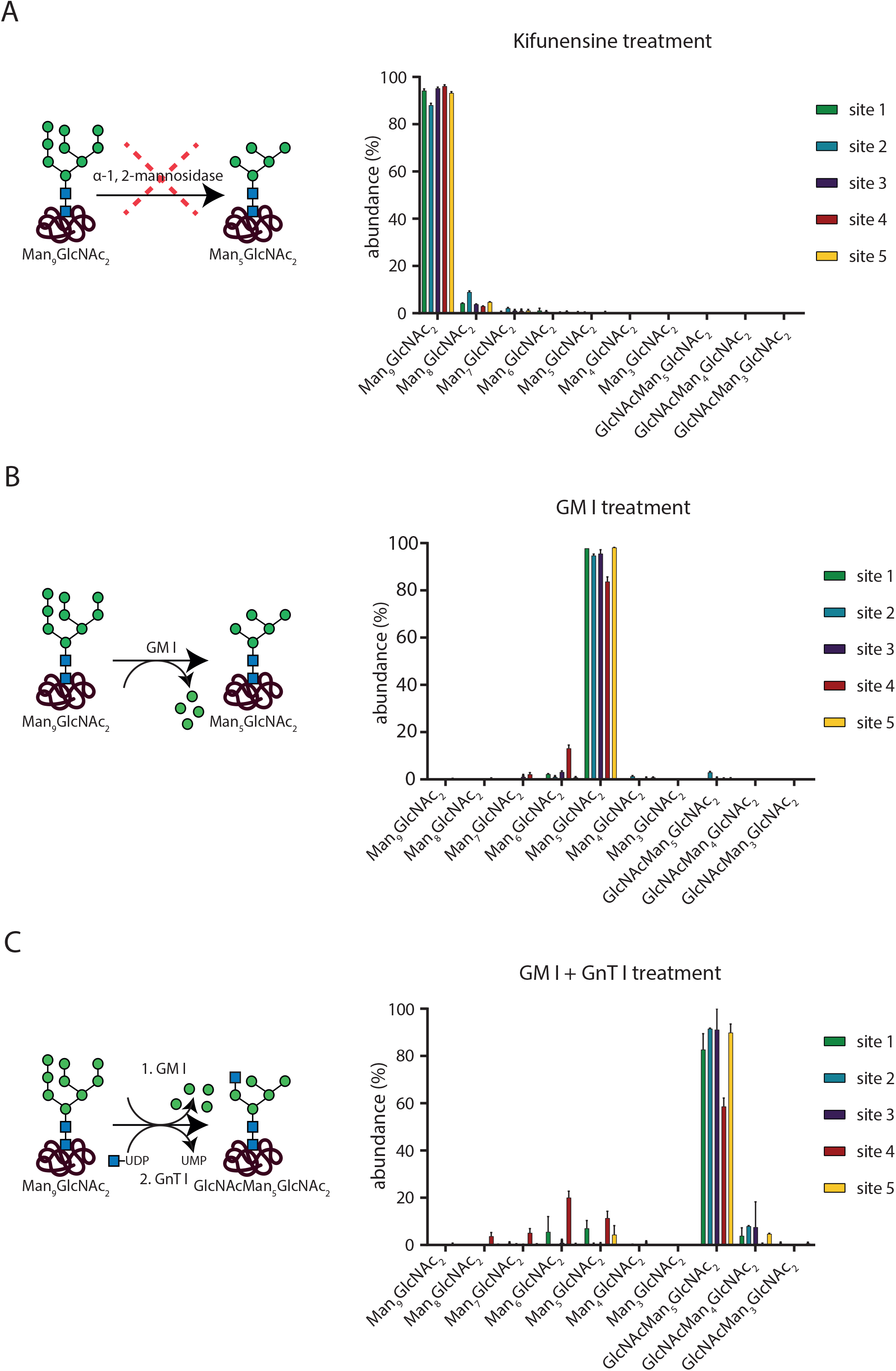
Generation of the PDI glycosubstrate for different glycan processing enzyme *in vitro* assays. The four different glycan processing enzymes used in the *in vitro* assays require different glyco-substrates. PDI was therefore treated according to these requirements. **A**: Substrate production for ER Man I and GM I *in vitro* assays. During PDI production in High-Five™ cells, the α-1, 2-mannosidase inhibitor kifunensine was added to the culture. It prevented mannose trimming in the ER and cis-Golgi. PDI purified from cells treated with kifunensine therefore showed a homogenous glycoform distribution (mainly Man_9_GlcNAc_2_ on all sites). **B:** Substrate production for GnT I *in vitro* assay. Purified PDI produced by High-Five™ cells not treated with kifunensine was incubated over night with GM I. Naturally occurring site-specific differences in glycoform distribution were adjusted by the mannose trimming activity of GM I, resulting in mainly Man_5_GlcNAc_2_ (the substrate of GnT I) on all sites. **C:** Substrate production for GM II *in vitro* assay. Purified PDI produced by High-Five™ cells not treated with kifunensine was incubated with GM I. After buffer exchange to GnT I activity buffer, PDI mainly bearing Man_5_GlcNAc_2_ on all sites was incubated with GnT I to obtain the glyco-substrate of GM II, GlcNAcMan_5_GlcNAc_2_.

**Figure EV 4:**
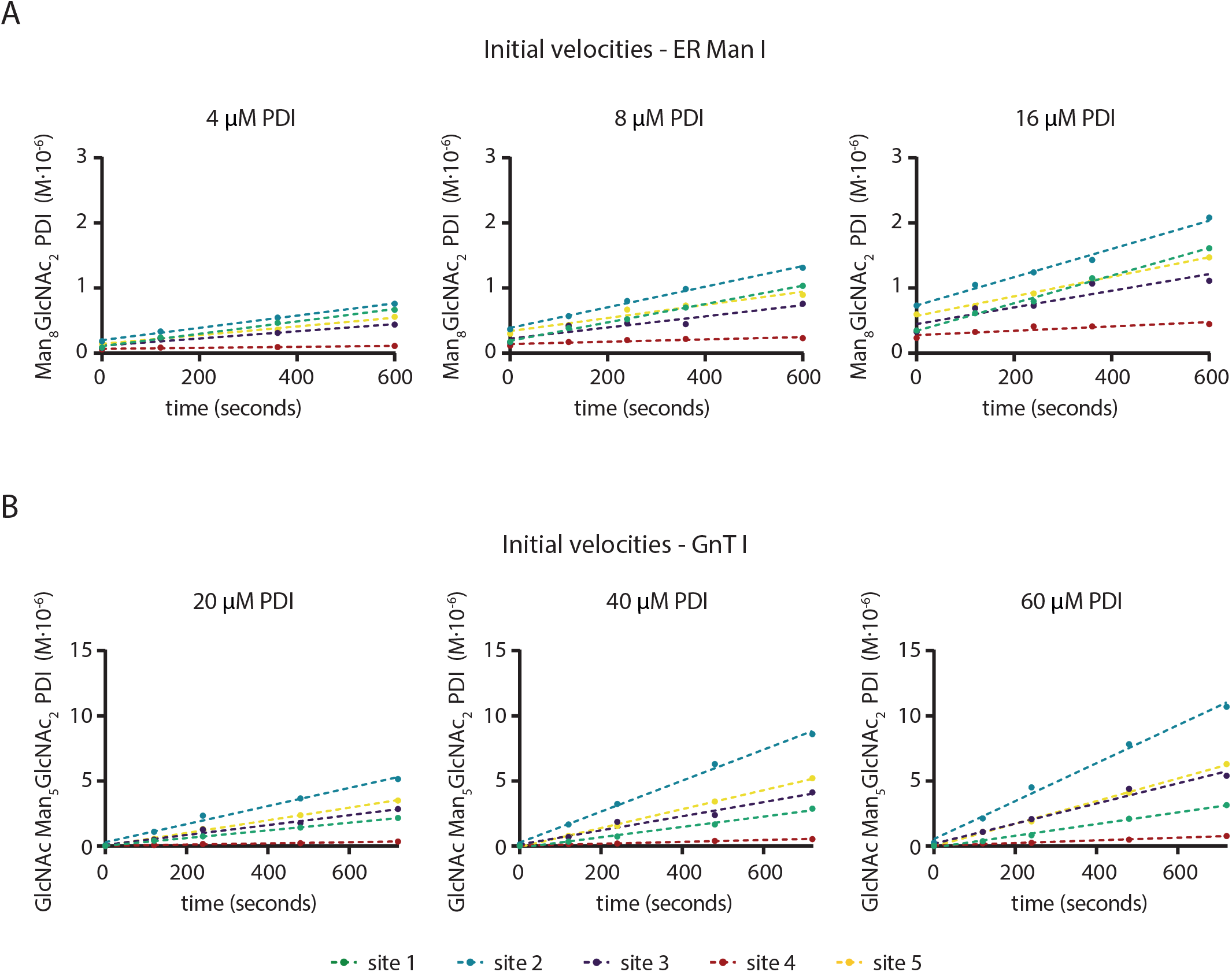
Linear phase of reaction for ER Man I and GnT I and different initial concentrations of PDI. **A**: ER Man I: The substrate Man_9_GlcNAc_2_-PDI (4, 8 and 16 μM) was incubated in presence of 0.6 nM ER Man I. The generation of the product Man_8_GlcNAc_2_ (in 10^−6^ M) on the five different sites is plotted against the incubation time (*n*=1). Linear regression lines were fitted using GraphPad Prism software. **B**: GnT I: The substrate Man_5_GlcNAc_2_-PDI (20, 40 and 60 μM) was incubated in presence of 67 nM GnT I and 5 mM UDP-GlcNAc. The generation of the product GlcNAcMan_5_GlcNAc_2_ (in 10^−6^ M) on the five different sites is plotted against the incubation time (*n*=1).

---

**Table EV 1.** Site specific choice of neighboring glycan and protein residues for the featurization step. Column one labels the considered site/glycan. The second column lists the neighboring glycans and the third column lists the residue indices of amino acids that are considered for the inverse distance featurization.

| Site | Glycan – Glycan | Glycan – Amino Acid |
| --- | --- | --- |
| 1 | 2, 3, 4 | 32 – 52, 87 – 107, 112 – 162 |
| 2 | 1, 4 | 32 – 52, 87 – 107, 112 – 126, 167 – 186, 218 – 232, 245 – 255, 317 – 327 |
| 3 | 1, 4 | 32 – 52, 87 – 107, 127 – 162, 187 – 192, 268 – 285, 345 – 355 |
| 4 | 1, 2, 3 | 32 – 52, 87 – 107, 112 – 192, 218 – 232 |
| 5 |  | 405 – 429, 483 – 504 |

**Table EV 2.** Amino-acid specific side-chain atoms that were used in hexose-amino acid distance calculations. The first column lists the amino acids and the second column lists the respective side-chain atoms, which were taken for the inverse distance calculation to respective O_5_ hexose atoms.

| Amino Acid | Side-Chain Atom |
| --- | --- |
| Arginine (ARG) | $C_\zeta$ |
| Asparagine (ASN) | $C_\gamma$ |
| Aspartic Acid (ASP) | $C_\gamma$ |
| Glutamine (GLN) | $C_\delta$ |
| Glutamic Acid (GLU) | $C_\delta$ |
| Histidine (HID) | $C_{\epsilon 1}$ |
| Lysine (LYS) | $N_\zeta$ |
| Serine (SER) | $O_\gamma$ |
| Threonine (THR) | $O_{\gamma 1}$ |
| Tryptophan (TRP) | $N_{\epsilon 1}$ |
| Tyrosine (TYR) | $O_\eta$ |

**Table EV 3.**
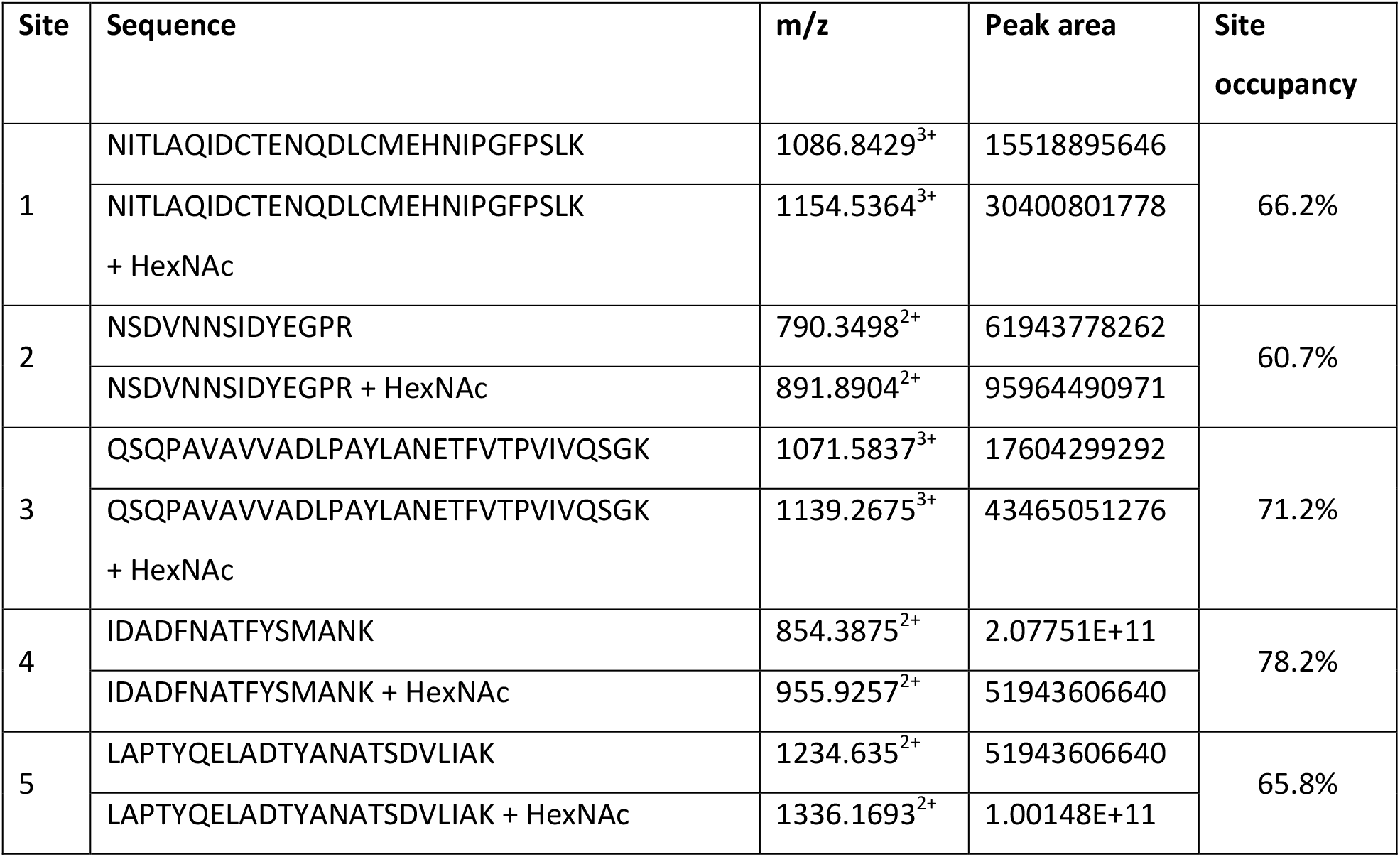
Glycosite occupancy of PDI produced in High-Five™ cells using the insect cell/baculovirus system.

| Site | Sequence | m/z | Peak area | Site occupancy |
| --- | --- | --- | --- | --- |
| 1 | NITLAQIDCTENQDLCMEHNIPGFPSLK | 1086.8429<sup>3+</sup> | 15518895646 | 66.2% |
|  | NITLAQIDCTENQDLCMEHNIPGFPSLK + HexNAc | 1154.5364<sup>3+</sup> | 30400801778 |  |
| 2 | NSDVNNSIDYEGPR | 790.3498<sup>2+</sup> | 61943778262 | 60.7% |
|  | NSDVNNSIDYEGPR + HexNAc | 891.8904<sup>2+</sup> | 95964490971 |  |
| 3 | QSQPAVAVVADLPAYLANETFVTPVIVQSGK | 1071.5837<sup>3+</sup> | 17604299292 | 71.2% |
|  | QSQPAVAVVADLPAYLANETFVTPVIVQSGK + HexNAc | 1139.2675<sup>3+</sup> | 43465051276 |  |
| 4 | IDADFNATFYSMANK | 854.3875<sup>2+</sup> | 2.07751E+11 | 78.2% |
|  | IDADFNATFYSMANK + HexNAc | 955.9257<sup>2+</sup> | 51943606640 |  |
| 5 | LAPTYQELADTYANATSDVLIK | 1234.635<sup>2+</sup> | 51943606640 | 65.8% |
|  | LAPTYQELADTYANATSDVLIK + HexNAc | 1336.1693<sup>2+</sup> | 1.00148E+11 |  |

**Table EV 4.** Site-specific initial ER Man I velocities (in 10^−9^ M/s ± SEM) for different substrate (PDI) concentrations.

|  | site 1 | site 2 | site 3 | site 4 | site 5 |
| --- | --- | --- | --- | --- | --- |
| 4 μM PDI | 0.95 ± 0.05 | 0.94 ± 0.04 | 0.55 ± 0.08 | 0.08 ± 0.02 | 0.67 ± 0.04 |
| 8 μM PDI | 1.41 ± 0.04 | 1.58 ± 0.07 | 0.85 ± 0.18 | 0.18 ± 0.05 | 1.01 ± 0.14 |
| 16 μM PDI | 2.12 ± 0.09 | 2.17 ± 0.14 | 1.28 ± 0.31 | 0.33 ± 0.1 | 1.5 ± 0.1 |

**Table EV 5.** Site-specific initial GnT I velocities (in 10^−9^ M/s ± SEM) for different substrate (PDI) concentrations.

|  | site 1 | site 2 | site 3 | site 4 | site 5 |
| --- | --- | --- | --- | --- | --- |
| 20 μM PDI | 2.95 ± 0.04 | 6.94 ± 0.46 | 3.80 ± 0.32 | 0.45 ± 0.05 | 4.9 ± 0.1 |
| 40 μM PDI | 3.98 ± 0.25 | 11.9 ± 0.44 | 5.43 ± 0.57 | 0.72 ± 0.06 | 7.26 ± 0.16 |
| 60 μM PDI | 4.5 ± 0.2 | 14.57 ± 0.72 | 7.72 ± 0.64 | 1.03 ± 0.07 | 8.69 ± 0.21 |

**Table EV 6.** Apparent rate constants k_1_, k_2_, k_3_ and k_4_ for formation of the Man_8_GlcNAc_2_, Man_7_GlcNAc_2_, Man_6_GlcNAc_2_ and Man_5_GlcNAc_2_ glycoforms by GM I on each glycosylation site, respectively.

|  | site 1 | site 2 | site 3 | site 4 | site 5 |
| --- | --- | --- | --- | --- | --- |
| $k_1$ ($\text{min}^{-1}$) | $0.21 \pm 0.03$ | $0.43 \pm 0.18$ | $0.37 \pm 0.08$ | $0.19 \pm 0.02$ | $0.45 \pm 0.16$ |
| $k_2$ ($\text{min}^{-1}$) | $0.22 \pm 0.04$ | $0.57 \pm 0.28$ | $0.26 \pm 0.05$ | $(2.7 \pm 0.3) \cdot 10^{-2}$ | $0.55 \pm 0.21$ |
| $k_3$ ($\text{min}^{-1}$) | $0.45 \pm 0.16$ | $1.10 \pm 1.0$ | $0.59 \pm 0.20$ | $(4.1 \pm 0.7) \cdot 10^{-2}$ | $1.10 \pm 0.7$ |
| $k_4$ ($\text{min}^{-1}$) | $(7.0 \pm 0.7) \cdot 10^{-3}$ | $(2.4 \pm 0.4) \cdot 10^{-2}$ | $(7.6 \pm 0.7) \cdot 10^{-3}$ | $(1.7 \pm 0.4) \cdot 10^{-3}$ | $(2.7 \pm 0.4) \cdot 10^{-2}$ |

**Table EV 7.**
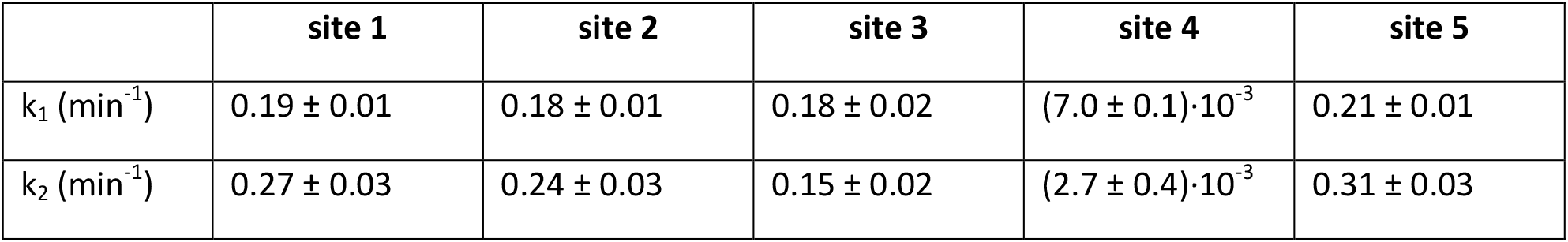
Apparent rate constants k_1_ and k_2_ for formation of the GlcNAcMan_4_GlcNAc_2_ and GlcNAcMan_3_GlcNAc_2_ glycoforms by GM II on each glycosylation site, respectively.

